# Similar mechanisms of retron-mediated anti-phage defense for different families of tailed phages

**DOI:** 10.1101/2024.02.09.579579

**Authors:** Fernando Manuel García-Rodríguez, Francisco Martínez-Abarca, Max E. Wilkinson, Nicolás Toro

## Abstract

Bacterial retrons are tripartite systems consisting of a cognate non-coding RNA, reverse transcriptase, and additional (effector) protein with diverse predicted enzymatic functions. In this study, we investigated the role and mechanism of Retron-Eco11, a novel type III-A3 retron system associated with a phosphoribosyltransferase-like effector protein, in phage defense. Here, we show that the Retron-Eco11 tripartite system protects against phage infection and that UvsW and D10, two functional homolog helicases found in T4 and T5 phages, respectively, serve as specific triggers for the Retron-Eco11 defense system. Our findings confirmed that msDNA and both protein components of the intron complex are indispensable for its protective function. Once the retron system detects the activity of these helicase proteins, it activates the toxicity of the effector protein bound to the retron complex, leading to an abortive infection. These findings underscore the application of a comparable anti-phage defense strategy using Retron-Eco11 across diverse phage families. This should aid in deciphering the processes through which the Retron complex detects and identifies the invading phages.

## Introduction

Bacterial retrons are tripartite systems composed of a cognate non-coding (nc)RNA, a reverse transcriptase (RT), and an additional (effector) protein or an RT-fused domain with an extraordinarily diverse range of predicted enzymatic functions^1–5^. Retrons are currently classified into 13 types on the basis of these three components, including additional variants^3^. Retron RTs use part of the ncRNA (msdRNA) as a template for synthesizing multicopy single-stranded DNA (msDNA), typically forming a DNA-RNA hybrid containing a 2′-5′ phosphodiester bond between the branching G of its RNA (msrRNA) component and the first nucleotide of the cDNA^6–10^.

Retrons were discovered more than 40 years ago, but their biological function remained elusive until recently. Retrons have been reported to function as anti-phage defense systems^1,2^ with an abortive infection (Abi) strategy. In addition, at least some retrons encode tripartite DNA-regulated toxin-antitoxin systems in which the RT and msDNA form an antitoxin unit that binds directly to the effector-associated protein, preventing its toxicity^4^. The retron defense system recognizes phage infection, possibly through the DNA stem loop of the msDNA^5^, triggering activation of the effector protein via an unknown mechanism. The recently determined cryo-electron microscopy (cryo-EM) structure of Ec86 (Retron-Eco1: type II-A3, the first retron discovered in *Escherichia coli*)^5^ has provided direct evidence that the effector protein stably associates with the other two components (msDNA and RT) of the retron complex, forming what appears to be a supramolecular assembly known as the retron-effector filament^11^.

Despite recent advances in retron biology, the mechanisms by which the retron complex senses and recognizes invading phages, modifying the retron-associated effectors, remain to be elucidated. We investigated a novel type III-A3 retron system known as Retron-Eco11, which is associated with a phosphoribosyltransferase (PRTase)-like effector protein. Our main objective was to explore the role and mechanism of the Retron-Eco11 system in the context of anti-phage defense. We identified the structural and functional counterparts of UvsW and D10, two helicases found in phages T4 and T5, respectively, as specific triggers of the Retron-Eco11 system. Once the retron system detects the activity of these helicase proteins, it activates the toxicity of the effector protein bound to the retron complex, leading to an abortive infection strategy. These findings highlight the use of a similar anti-phage defense mechanism by Retron-Eco11 for different phage types and should help unravel the mechanisms by which the retron complex senses and recognizes invading phages, triggering the abortive infection process.

## Results

### The Retron-Eco11 tripartite system

The Retron-Eco11 system was previously identified^3^ in the whole-shotgun sequence of *Escherichia coli* EC2608. Eco11 is a tripartite system comprising a reverse transcriptase from clade 9, a non-coding RNA gene transcribed as a retron RNA with a predicted conserved type IIIA structure, and a gene encoding an effector protein grouped within protein cluster 67_3 (**Figure 1A**). This effector protein is characterized by an N-terminal phosphoribosyltransferase (PRTase)-like domain and a C-terminal RuvB winged helix DNA-binding domain.

**Figure 1.**
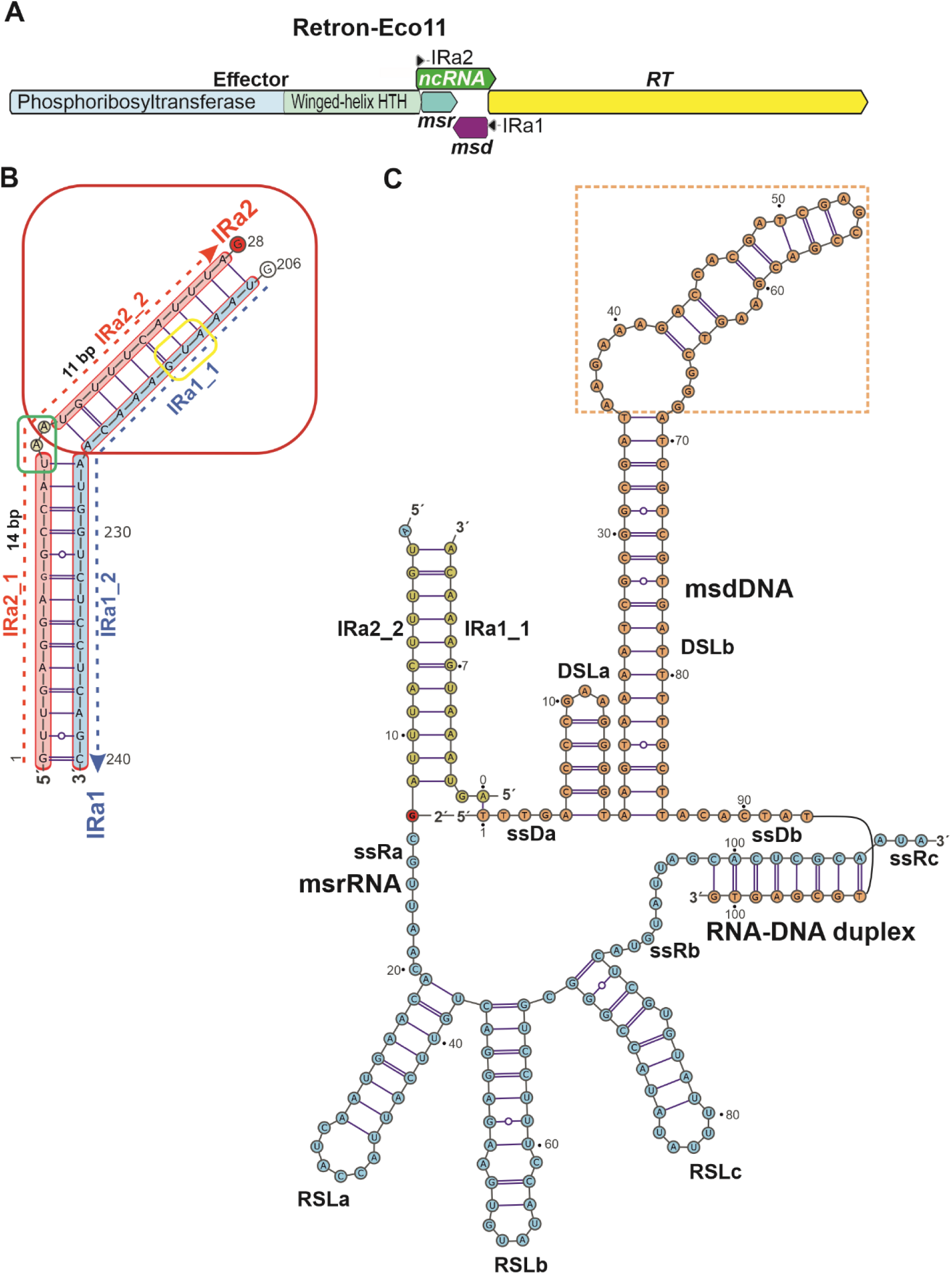
Retron-Eco11 tripartite system. **(A)** Schematic diagram of the Retron-Eco11 system in the *E. coli* (EC2608) genome. **(B)** Schematic representation of the predicted pair of inverted repeats msrRNA and msdRNA (IRa2 and IRa1) located at the 5′ and 3′ ends of the msrRNA and msdRNA, respectively. The stop codon of the effector protein that include two bulged adenosines at the 5′ side of the msrRNA transcript, and the translational start codon of the RT are framed by green and yellow rounded rectangles, respectively. The predicted branching G (msrRNA rG28) marked by the red circle and the msdDNA dT1 are predicted to form a 2′-5′-phosphodiester bond. **(C)** Predicted secondary structure of the Retron-Eco11 msDNA, according to the model reported for EC86^5^. The msdDNA, msrRNA, single-stranded (ss) DNA, and RNA regions, and the DNA-RNA duplex, are indicated. The predicted (**Figure S1**) flexible DSLb hairpin loop is framed by a dashed-line rectangle.

The Eco11 ncRNA overlaps the 3′ and 5′ ends of the effector and RT genes on the same strand. It contains two regions, the msrRNA and msdRNA regions (**Figure 1B**), each with a pair of inverted repeats (IRa2 and IRa1) located at the 5′ and 3′ ends of msrRNA and msdRNA, respectively. This pairing involves 25 base pairs, which is unusually long for retrons, as typical IRs range from 8 to 15 base pairs. It includes an outer arm of 14 base pairs (IRa2_1/IRa1_2) and an inner arm of 11 base pairs (IRa2-2/IRa1_1), separated by two bulged adenosine residues on the 5’ side of the msrRNA transcript. The inner arm of 11 base pairs (IRa2-2/IRa1_1) is sufficient for the generation of msDNA, which is crucial for the biological activity of the retron system.

The IR region is followed by unpaired G residues, and the msrRNA rG28 (branching G) and msdDNA dT1 are predicted to form a 2′,5′-phosphodiester bond. Apart from the single-stranded RNA segments, the predicted msrRNA region, which is not reverse-transcribed, comprises three short RNA stem-loops and one DNA-RNA duplex. According to the predicted structure and the identified 3′ end of the msdDNA, the RT-Eco11 appears to produce a 101-base msdDNA that is predicted to fold into two DNA stem-loop structures, DSLa and DSLb, comprising 13 and 69 nucleotides, respectively. The predicted secondary structure includes a stable conserved 18-base pair stem at the 5’ end of DSLb, with the hairpin loop displaying greater structural flexibility (**Figure 1C**; **Figure S1**). These findings are consistent with previous reports suggesting that a segment of the long stem-loop in msDNA can serve as a template for precise genome editing through homology-directed repair (HDR)^12,13^.

As anticipated, the sequences of RT-Eco11 and RT-Ec86 (RT-Eco1), both of which belong to clade 9, were well aligned. They displayed a pairwise identity of 24.5%, encompassing the RT1-7 regions and the retron-specific regions, denoted X and Y. In these retron-specific regions, the conserved NAxxH motif was replaced by AAxxH, and the VTG motif was replaced by VLG (**Figure S2A**). Furthermore, a superimposition of the predicted 3D structures of RT-Eco11 generated by AlphaFold2 (**Figure S2B**) and RT-Eco1 associated with its cognate msDNA (Ec86 complex, Protein Data Bank: 9V9U) gave a root mean square deviation (RMSD) of 2.55 over 320 Cα atoms, indicating structural similarities between these two molecules.

The identified PRTase-like effectors associated with retrons are closely related phytogenetically and distinctive of retron type III systems^3^. In these systems, the PRTase-like effector may occur alone or be followed by a downstream-encoded protein with a predicted winged-helix DNA-binding domain (**Figure 2A**). In certain variants of type III systems, such as A3, A4, and A5, this DNA-binding domain is fused to the C-terminus of the putative PRTase. Sequence comparisons revealed that proteins within cluster 67-3 possess a short conserved (24 amino acid residues) tail, referred to as the C-terminal tail (CTT). This CTT can be divided into two regions: a hypervariable C-terminal linker (CTL, spanning residues 390-404 in Eco11) and a conserved C-terminal peptide **YYP**xx**LR**xx (CTP, residues 405-413 in Retron-Eco11) (**Figure 2A, 2B**).

**Figure 2.**
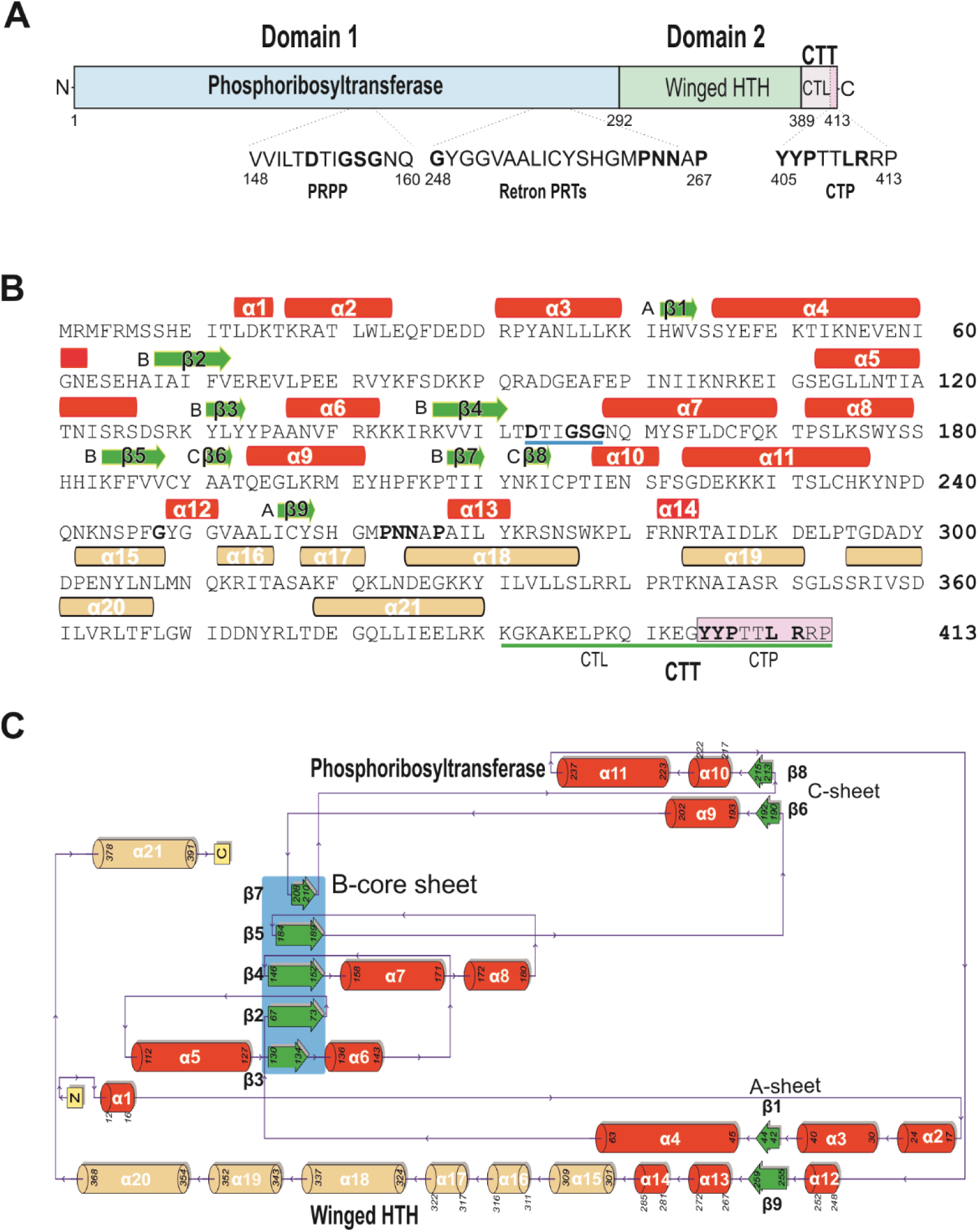
Retron-Ec11 phosphoribosyltransferase effector protein. **(A)** Protein domains of the Ec11-associated phosphoribosyltransferase. The figure highlights the PRPP motif, the retron-specific motif, and the C-terminal tail (CTT). It also includes the C-terminal linker (CTL) and the characteristic C-terminal peptide (CTP). Residues conserved within phosphoribosyltransferase protein cluster 67-3^3^ at an identity threshold of 90% are indicated in bold. **(B)** Sequence and secondary structure of the Ec11-effector protein. The alpha helices within the phosphoribosyltransferase and winged-HTH domains are colored in red and brown, respectively. The PRPP binding site is marked by a blue line and conserved residues within the characteristic motifs are shown in bold. The CTT is identified by a green line below the sequence, and the CTP is boxed. **(C)** Schematic diagram illustrating the structure of the Ec11 PRTase-like effector protein, as obtained from PDBsum^33^. The characteristic B-core sheet of PRTases (comprising β3, β2, β4, β5, β7) sandwiched between α6, α7, and α8 on one side of the β-sheet, with α5 on the other side, is highlighted in blue.

Type I PRTases serve as both enzymes and regulatory proteins in nucleotide synthesis and salvage pathways. They can be identified by a conserved PRPP (5-phospho-α-D-ribose-1-diphosphate) binding motif and a common core structure featuring five-stranded, parallel β-sheets surrounded by three or four α-helices, a structural arrangement closely resembling a typical Rossmann dinucleotide binding fold^14^. The predicted structure of the Eco11 effector includes a characteristic B-core sheet of PRTases (comprising β3, β2, β4, β5, β7) sandwiched between α6, α7, and α8 on one face of the β-sheet, with α5 on the other face (**Figure 2B, 2C**). The Eco11 effector carries the VVIL**<u>TD</u>**<u>TI**GSG**</u>NQ (residues 148-160) PRPP motif, in which the PRPP binding site (underlined) is located in the PRPP loop between β4 and α7. In this subdomain, PRTase-like proteins associated with retrons contain a distinctive conserved motif **G**x14**PNN**x**P**^3^ in the loops flanking the β9 strand of the Eco11 effector (**Figure 2A, 2B**). This motif may reflect a common biochemical function of these proteins.

A FATCAT-search for similar protein structures^15^ revealed that the protein structures most similar to the Eco11 effector (**Figure S3A**, **S3B**) were those of xanthine-guanine-hypoxanthine PRTase (with an RMSD of 3.07 over 130 Cα atoms) and the purine PRTase (with an RMSD of 3.09 over 132 Cα atoms). Phylogenetic analyses provided further evidence that the retron-associated effector in cluster 67-3 represents a distinctive lineage, probably constituting a novel and previously unreported group within the type I PRTases (**Figure 3**). The precise role or function of this particular group remains undetermined and warrants further investigation.

**Figure 3.**
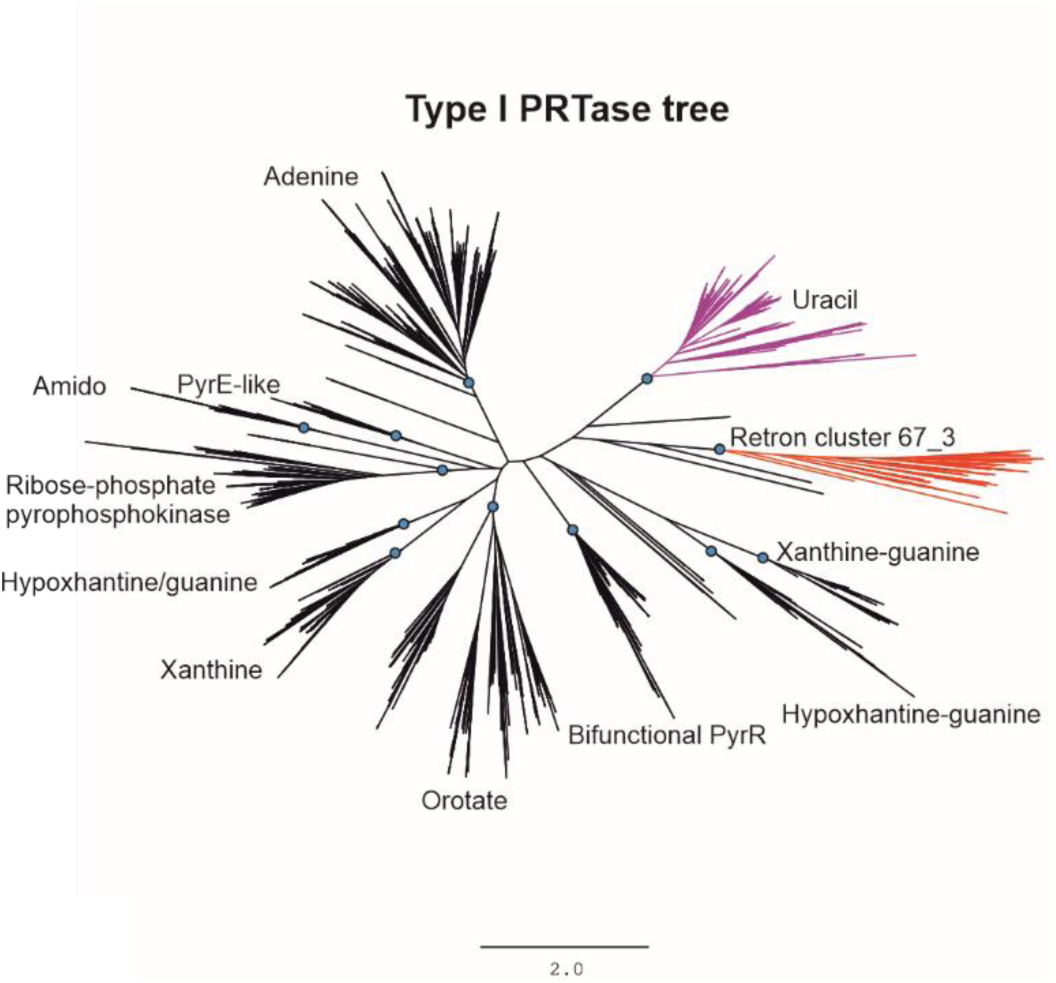
Phylogeny of type I PRTases and retron type III_A-associated phosphoribosyltransferases. The unrooted tree, constructed from an alignment of 2564 sequences carrying the phosphoribosyltransferase domain (IPR000836) and 127 sequences from the retron type III-A associated protein cluster 67_3^3^ was generated with FastTree. In this tree, the cluster containing 67_3 sequences is shown in red, whereas a nearby clade of uracil phosphoribosyltransferases is indicated in purple for reference.

The PRTase-like domain of Eco11 is followed by a C-terminal winged helix-turn-helix (HTH) domain, the predicted 3D structure of which aligns with transcriptional regulators, with an RMSD of 2.00 over 72 Cα atoms (**Figure S3C**). Interestingly, in the predicted 3D structure of the Eco11 effector, the C-terminus appears as a disordered coil-coil region. When the C-terminal winged HTH domain of the Eco11 effector was superimposed on that of the retron Ec86 effector, the RMSD was 2.55 over 73 equivalent positions (**Figure S3C**). The C-terminus of the Ec86 effector also has a structurally similar terminal peptide, flanked by tyrosine (residue 299) and proline (residue 307) residues. This terminal peptide is required for effective phage defense and appears to interact with its cognate msrRNA and msdDNA^5^. Thus, the C-terminal tail of the Eco11 effector may be available for binding to and modifications by other proteins, as suggested for the Ec86 effector.

### The Retron-Eco11 tripartite system protects against phage infection

We studied the function of Retron-Eco11 and the mechanism by which it acts as an anti-phage defense system, by synthesizing the retron system of EC2608 and expressing it in the *E. coli* MG1655 strain, which lacks retrons. Retron-containing bacteria were tested against a panel of 12 phages. Significant anti-phage activity was observed against *E. coli* phages T2, T4 (*Straboviridae*) and T5 (*Demerecviridae*), and a decrease in plaque size was observed for phage T7(*Autographviridae*) (**Figure 4**).

**Figure 4.**
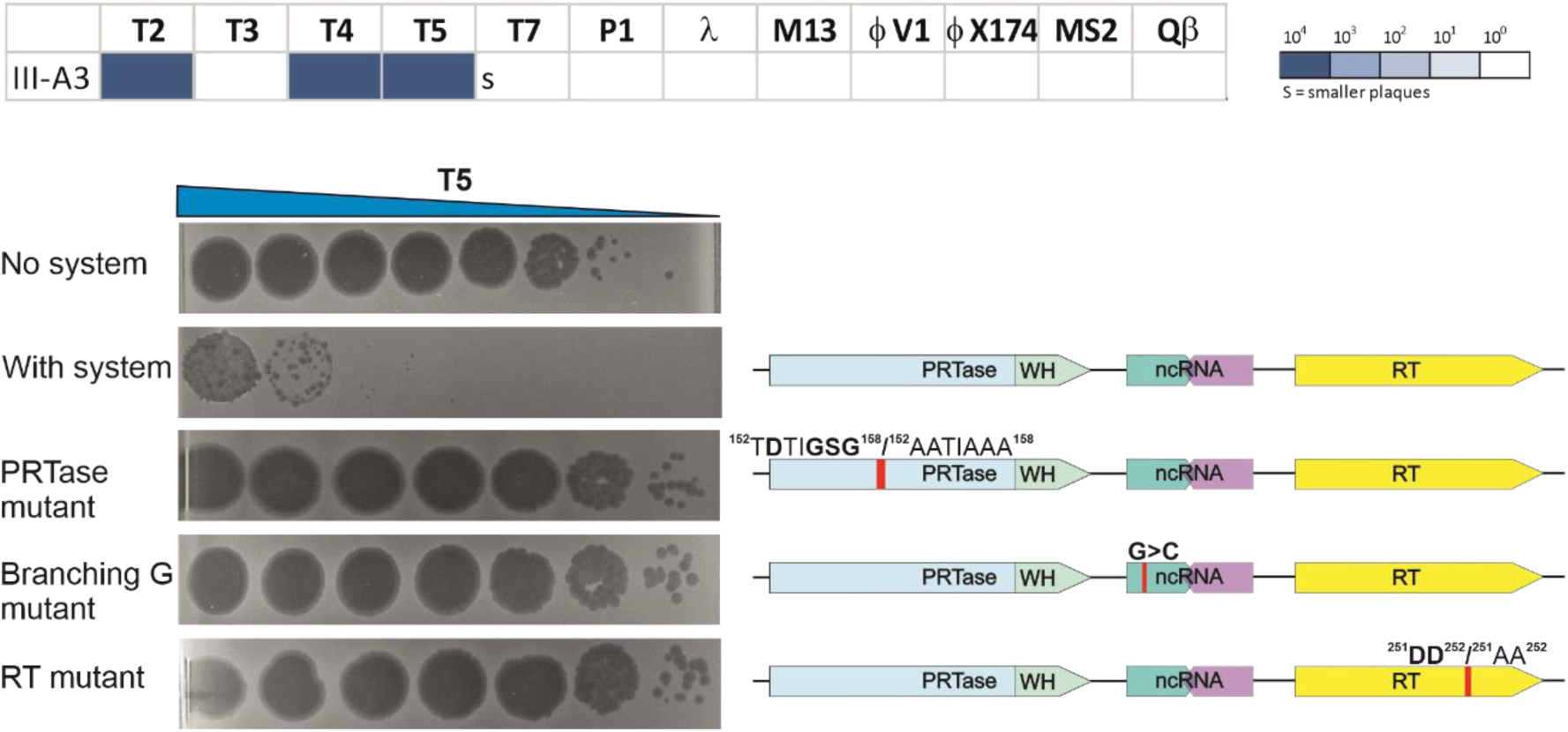
Protection against phage infection mediated by the Retron-Eco11 system. Fold-protection was determined in serial-dilution plaque assays comparing efficacy between the system-containing strain and the control strain lacking the retron system. The data displayed are representative results from three replicates. In the case of the T7 phage, a significant decrease in plaque size was observed, indicated with an “S” for smaller plaques. The phages used in these assays are indicated. For the sake of simplicity, only T5 plaques are shown here. On the right-hand side, schematic representations are provided for the wild-type and mutant constructs for each component of the system used in the assays.

We investigated the components of the system essential for this observed defense activity by performing assays on mutants with modifications to the catalytic sites of the effector protein (152T**D**TI**GSG**158/152AATIAAA158), the RT (**D**251A/**D**252A) and the predicted branching **G** (changed to C) of the ncRNA. All of these mutations abolished the ability of Retron-Eco11 to protect against phage infection. These findings confirm that the msDNA and both the protein components are indispensable for the protective function of this retron system.

### Different bacteriophages trigger the Retron-Eco11 defense system via a common mechanism

T4 and T5 are bacteriophages with contrasting genome sizes, genetic contents, and host ranges^16–20^. We investigated the ways in which Retron-Eco11 specifically recognizes and responds to phage infections, by conducting experiments to isolate phage T4 and T5 mutants capable of evading this defense system. For each phage, we successfully isolated five mutants capable of disabling retron defense activity.

All the T4 mutant phages presented two deletions of different sizes in the UvsW RNA-DNA-encoded helicase^21–24^. These deletions led to frameshift mutations resulting in prematurely truncated N-terminal RecA-like motor domain proteins. The T5 mutants displayed a different pattern. They had single point mutations leading to the replacement of individual amino-acid residues in the C-terminal RecA-like domain of the D10 helicase^25^ (**Figure 5A**). D10 is a functional homolog of UvsW, and both proteins belong to the SF2 helicase family^26,27^.

**Figure 5.**
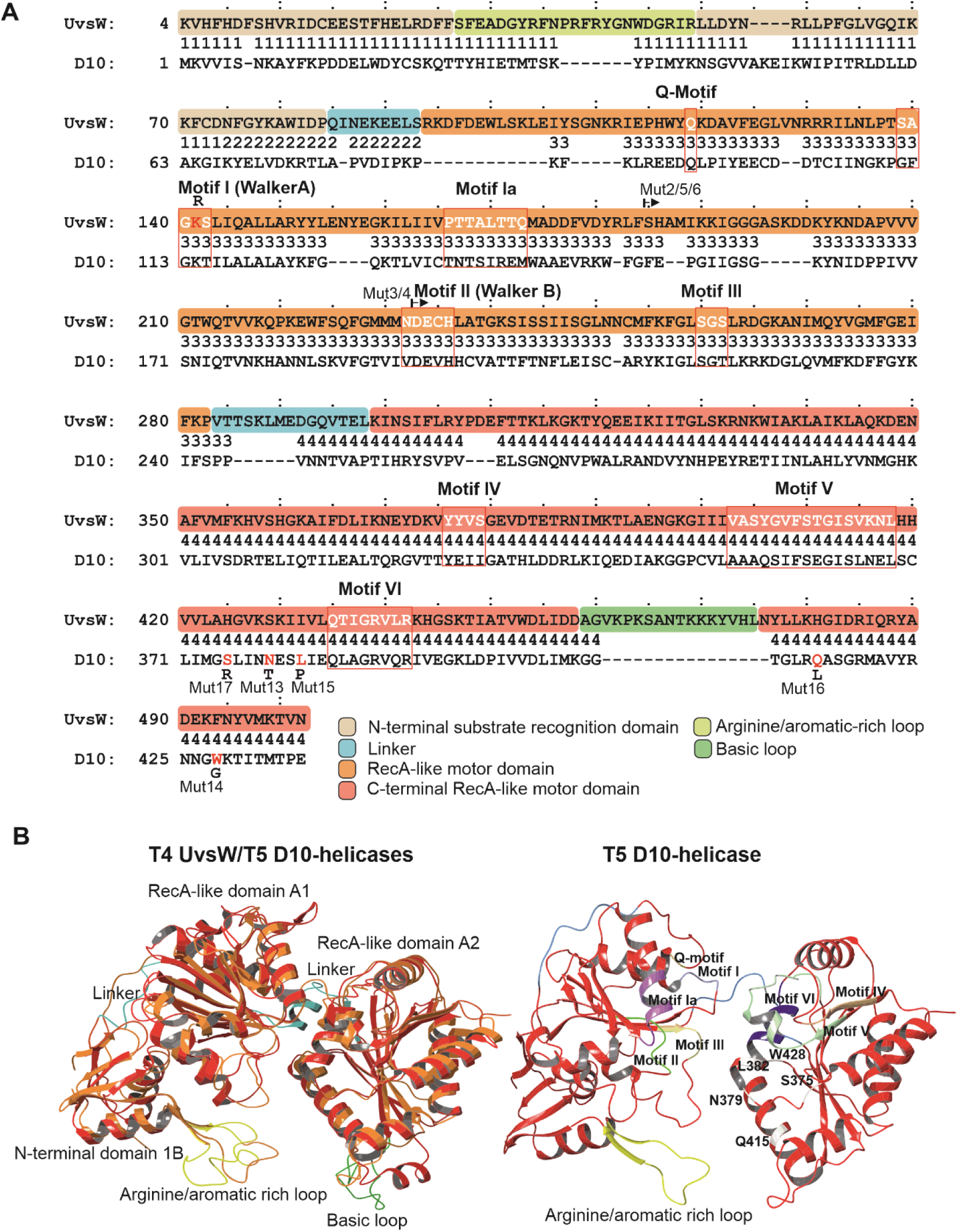
Structural similarities between the T4 phage UvsW and T5 phage D10 helicases. (**A**) FATCAT-pairwise alignment^15^ of T4 phage UvsW helicase (PDB: 2OCA) and the predicted D10 helicase generated with AlphaFold2. FATCAT was used for flexible structural alignment with the chaining of aligned fragment pairs to allow twists. The numbers between alignments are the block index. The four-domain architecture of UvsW^34^ is indicated and highlighted in different colors. The Sf2 helicase motifs (I to VI) are boxed. The Q-motif, arginine/aromatic and basic loops are also shown The region missing from the UvsW truncated proteins of T4 phage escape mutants is indicated by small arrows, and single-point mutations leading to substitutions in the C-terminal RecA-like motor domain of the D10 protein of the T5 phage escape mutants are highlighted in red, with the corresponding substitutions indicated. (**B**) Superimposed structures of T5 phage D10 helicase (in red, obtained with AlphaFold2) and the T4 UvsW (in orange) helicase. The predicted structure of T5 D10 is shown in the right panel with the Q-and motifs I to VI, and the residues substituted in the phage escape mutants are indicated. There are 432 equivalent positions, with an RMSD of 2.78Å, and three twists.

We hypothesized that the truncated UvsW proteins, if produced, would have no helicase activity, as they would lack conserved sequence motifs II and VI, which are essential for ATP binding and hydrolysis, and motifs IV and V, which are crucial for binding to nucleic acids. However, as the crystal structure of the D10 helicase had not been determined, uncertainties remained concerning the impact on the biochemical properties of D10 of the single amino-acid substitutions found in the mutants. We used AlphaFold2 to predict the structure of D10 (**Figure 5B**), to shed light on the potential functions of these residues. The predicted structure of D10 matched that of UvsW over 432 equivalent positions, with a root mean square deviation (RMSD) of 2.78Å, indicating that these two proteins are structural homologs. The amino-acid substitutions in D10 (S375R, N379T, L382P, Q415L, and W428G) were located in the cleft facing the N-terminal conserved RecA-like helicase domain (**Figure 5B**). Most of the substituted amino acids were in close contact with either motif V or motif VI residues, a direct hydrogen bond being observed between residues L382 and L386 in motif VI (**Figure S4**). These structural findings are consistent with an abolition of the helicase activity of D10 by the identified mutations. Taken together, these results suggest that inactivation of the UvsW/D10 helicases played a crucial role in enabling T4 and T5 phages to evade the Retron-Eco11 defense system, possibly due to an impairment of the sensing or biochemical activities of these proteins.

### The UvsW and D10 helicases serve as triggers for the Retron-Eco11 defense system

The Retron-Eco11 system did not inhibit cell growth when expressed in *E. coli*. Furthermore, the effector protein, when expressed independently via an inducible arabinose promoter plasmid construct (pBAD), displayed no active toxicity (**Figure 6A**).

Additional experiments were performed to confirm that T4 and T5 phages activated the Retron-Eco11 defense system via a similar mechanism. These experiments involved the expression of wild-type and mutant UvsW/D10 proteins in *E. coli* cells, in both the presence and absence of the retron system and a mutant form of the effector protein. In the presence of the complete retron system, the expression of wild-type UvsW/D10 resulted in toxicity (**Figure 6B**). However, no such impairment of cell growth was observed when mutant helicases were expressed, in the absence of the retron system, or in the presence of a mutated effector protein. These findings provide further compelling evidence to support the notion that the UvsW/D10 helicases themselves serve as triggers for the Retron-Eco11 defense system. Moreover, they suggest that effector protein toxicity is induced by recognition of a phage trigger by the retron complex, supporting the notion that the Retron-Eco11 system responds to specific phage-related cues or activities.

**Figure 6.**
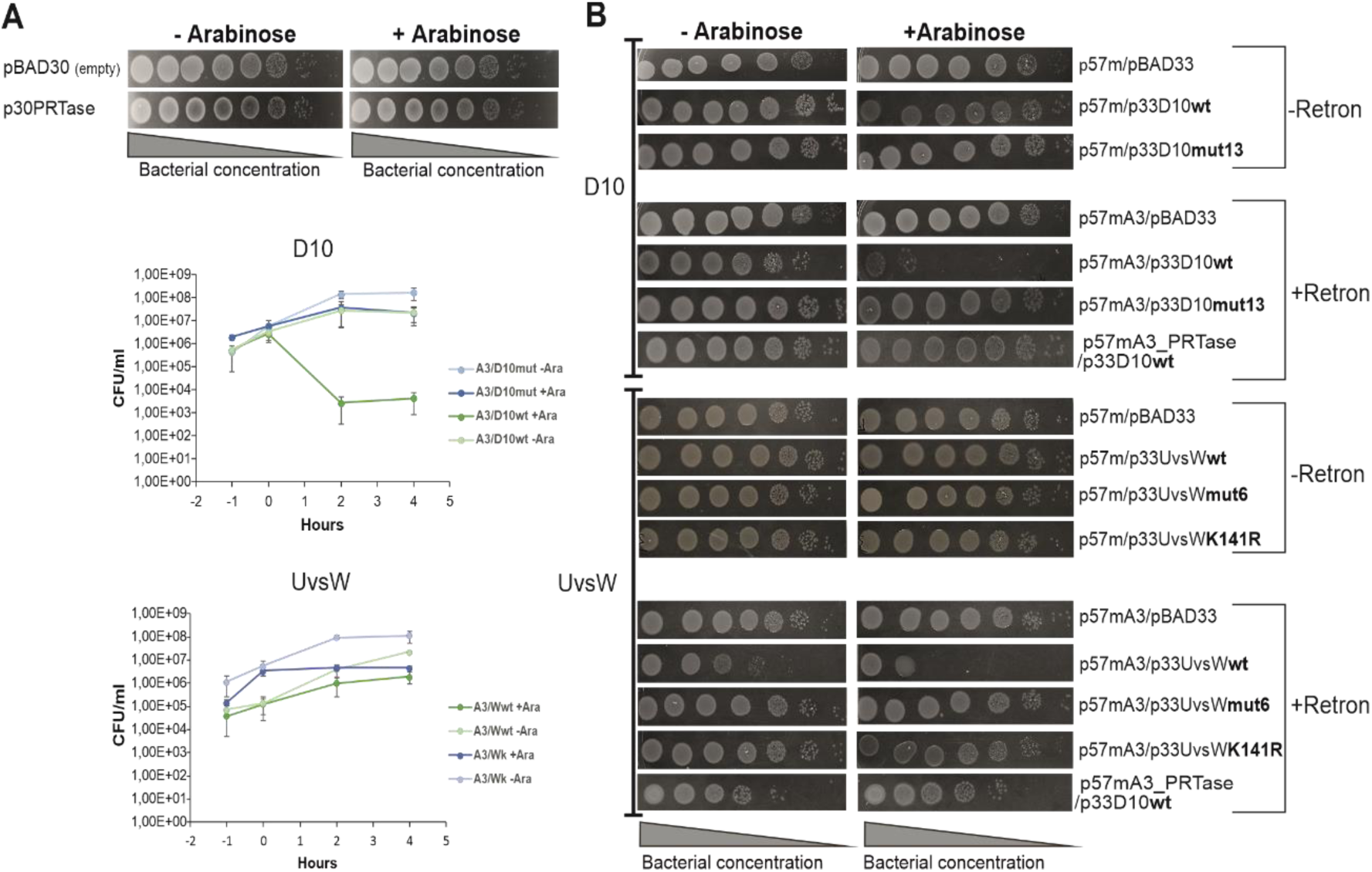
The UvsW and D10 helicases trigger the Retron-Eco11 defense system. **(A)** The PRTase effector is not toxic on its own in the *E. coli* host. Bacterial cell viability was assayed in the presence (p30PRTase) or absence (pBAD30) of the PRTase effector, expressed under the control of an arabinose-inducible promoter in pBAD. Growth curves for *E. coli* harboring the Retron-Eco11 defense system and expressing phage helicases D10 and UvsW, or their mutant derivatives, are also shown. These helicases were also encoded by pBAD and were expressed under the control of an arabinose-inducible promoter. A3 is p57mA3; D10 is p33D10wt; D10mut is p33D10mut13; Wwt, is p33UvsWwt; Wk is p33UvsWK141R. (**B**) Expression of the T4 UvsW and T5 D10 helicases results in toxicity in a strain containing the Retron-Eco11 system. Bacterial cell viability was measured in the presence (p57mA3) and absence (p57m) of the retron system and in the absence (pBAD33) and presence of the T4 and T5 genes encoding the UvsW and D10 helicases or their mutant derivatives. The PRTase mutant was also included in some of the assays. Cell viability was assessed by plating 10-fold serial dilutions of cells with (+Arabinose) and without (-Arabinose) induction of the phage genes.

We investigated the ability of the retron to recognize the presence of the helicases through protein-protein interactions or from their activity, by testing a mutant form of UvsW (K141R) with a substitution in the Walker A motif. This mutation is known to inhibit ATP hydrolysis significantly and to prevent translocation (branch migration), but it retains the ability to bind Holliday Junction (HJ)-containing structures^22^. Interestingly, the expression of this UvsW mutant helicase did not result in toxicity in cells containing the Retron-Eco11 system (**Figure 6B**). This observation suggests that the Retron-Eco11 defense system may respond to the translocation activity of these phage proteins rather than acting through a direct interaction between proteins. Furthermore, an assessment of bacterial viability after induction of the phage helicases D10 and UsvW and their mutant derivatives (**Figure 6A**) revealed a substantial, faster decrease in the number of colony-forming units (CFU) following D10 expression. This finding suggests that the D10 helicase may have a more pronounced role in activating the retron defense system.

### The Retron-Eco11 defense mechanism relies on specific features of the UvsW and D10 phage helicases

Our results, as presented above, suggest that Retron-Eco11 can detect the branch migration activity of UvsW/D10 phage helicases to some extent. The functions of the UvsW helicase are similar to those of the RecG protein, which is known to resolve stalled replication forks and to facilitate the branch migration of Holliday junctions. However, despite their functional similarities, these proteins display little significant sequence or structural similarity beyond the conserved motifs of SF2 helicases. Furthermore, RecG is a much larger molecule that requires additional domains for substrate recognition and binding^28^.

We investigated the possibility that the Retron-Eco11 defense mechanism monitors the functions common to these two helicases by performing testing whether the retron system in *E. coli* cells could recognize the absence of RecG. Interestingly, transformation with a Δ*recG* mutant in plasmids containing Retron-Eco11 was efficient, with no apparent growth arrest in the transformed cells. When co-expressed with UvsW/D10 helicases and their mutant derivatives, the phenotype resembled that of wild-type *E. coli* cells (**Figure S5**). These findings suggest that the Retron-Eco11 defense mechanism is not triggered exclusively by the activities common to the RecG and UvsW/D10 helicases. Instead, it appears to be specific to certain characteristics or features of these phage helicases that remain incompletely understood.

### Retron-Eco11 complex interactions

We investigated interactions within the retron complex by adding a 3xFlag tag to the C-terminus of RT-Eco11. This addition had no effect on the biological anti-phage defense activity of the complex. We then performed immunoprecipitation experiments to identify the components of the retron complex through mass spectrometry proteomics analysis. As expected, other than the RT, the only protein for which significant enrichment was observed was the PRTase effector protein. Furthermore, the total DNA extracted from these samples contained the matured msDNA. None of the retron components were detected in samples with the untagged RT, confirming the specificity of the interactions within the retron complex. However, when a retron construct with a mutation of the branching G of the ncRNA was used, no binding of the PRTase to the RT was detected. This observation indicates that, contrary to observations for other retrons^4^, the presence of the msDNA is essential for the effector protein to bind to the retron complex in the Retron-Eco11 system. Thus, the msDNA appears to play a crucial role in stabilizing the interaction between the RT and the effector protein within this particular retron system (**Figure 7A, 7B**).

**Figure 7.**
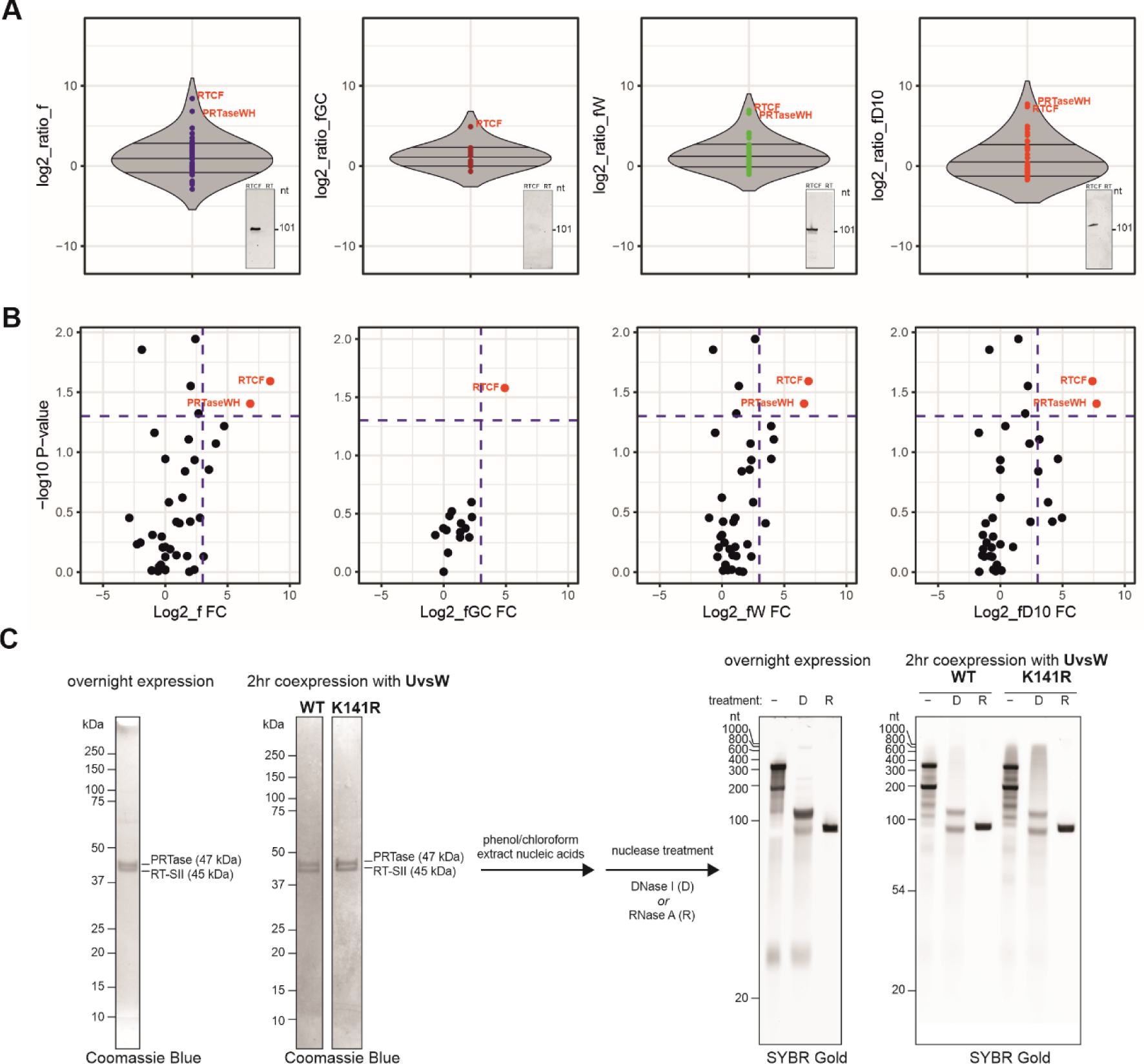
Formation of the Retron-Eco11 complex consisting of the three components of the tripartite system. (**A**) Violin plots are used to illustrate the protein enrichment achieved through co-immunoprecipitation with retron-RT-3xFlag. As shown in the log2_ratio_f panel, the components of the retron complex include the RT, the PRTase (PRTaseWH), and the retron msDNA (shown as an inset within the frames). RTCF is the RT tagged with a FLAG marker at its C-terminus. However, when a retron construct with a mutation of the branching G of the ncRNA was used, no binding of the PRTase to the RT was detected (log2_ratio_fGC). Prolonged incubation of the beads for over 24 hours with protein extracts containing the UvsW (log2_ratio_fW) or the D10 (log2_ratio_fD10) helicase did not affect the composition of the retron complex. The *y*-axis shows the mean log_2_ ratio for the identified proteins (detected with more than two peptides) for RT-3xFLAG relative to proteins identified with untagged RT as a control. Please note that the RT and PRTase were not detected in the control samples, and the corresponding missing values are replaced with the minimal emPAI value of the samples for the purpose of statistical analysis^35^. (**B**) Volcano plots displaying the results of affinity purification for the experiments detailed in panel A. The log_2_-fold protein enrichment (*x*-axis) is plotted against the corresponding *p*-value (-log_10_ *p* value). (**C**) Retron complex purification. Co-expression *in vivo* of the retron and the UvsW, UvsWK141R mutant helicases revealed no significantly alteration to the stoichiometry of the retron complex. The migration of the RNA and DNA components of the retron complex on a denaturing acrylamide gel was also unaffected by *in vivo* co-expression with UvsW or UvsWK141R

We also investigated whether the expression of the UvsW/D10 phage helicases affected the composition of the retron complex. The co-incubation *in vitro* of the immunoprecipitated retron complex with total proteins including the phage helicases (**Figure 7A, 7B**), and the co-expression *in vivo* of the retron and the UvsW, UvsWK141R mutant, and D10 helicases (not shown) revealed no significantly alteration to the stoichiometry of the retron complex. The migration of the RNA and DNA components of the retron complex on a denaturing acrylamide gel was also unaffected by *in vivo* co-expression with UvsW or UvsWK141R (**Figure 7C**). These findings suggest that the activation and toxicity of the effector protein probably occur within the complex itself, suggesting a central role of this complex in the recognition of phage infection and in modulating the activity of the effector protein.

## Discussion

In this study, we report a new retron system (Retron-Eco11: type III-A3), describe its function in anti-phage defense, and identify the specific phage helicases that activate the retron response. Retron-Eco11 protects *E. coli* cells against both T-even and T-uneven phages, apparently through a similar mechanism involving recognition of the expression of the homologous T4 UvsW and T5 D10 helicases. The effector of this retron system is a PRTase-like protein that is not toxic to the host cell on its own but becomes toxic when bound to the retron complex following the recognition of phage triggers, ultimately leading to abortive infection through cell growth arrest and death.

Despite recent advancements in retron biology, the precise mechanisms governing the ability of the retron complex to detect and respond to invading phages remain enigmatic, and many observations are puzzling. Several phage triggers activating retron defense systems have been identified in recent studies^29^. The inhibition of RecBCD by phage proteins has been reported to activate retron Ec48 (Retron-Eco6: type IV), leading to cell growth arrest and, ultimately, cell death^2^. A similar activation mechanism appears to occur for retron Sen72 (Retron-Sen1: type V), albeit involving a different effector protein. The encoded reverse transcriptases (RTs) of these retrons share a common phylogenetic origin, both belonging to clade 3. However, not all retron systems with RTs of identical phylogenetic origin follow the same activation pattern. For instance, Retron-Eco8 (type I-B2), which also belongs to clade 3, is activated by the expression of single-stranded binding (SSB) proteins from phages of the *Podoviridae* family^29^. Furthermore, different phage proteins are responsible for triggering different retron defense systems. In the case of retron Ec67 (Retron-Eco2: type I-C1), which has a clade 8 RT, activation appears to be associated with the early-expressed phage proteins A1 and DenB, both of which are involved in DNA degradation^29^. Similarly, the toxicity of retron Ec86 (Retron-Eco1: type A3) seems to be triggered by the phage protein Dcm, which modifies the msDNA^11^. Thus, even though the mechanism of phage recognition is probably common to different retrons, each component of a retron system and the various combinations of components may contribute to the establishment of a different model of defense activity.

Retron Ec86 and Retron-Eco11 both have clade 9 RTs, but their msDNA components have different structural characteristics and are associated with different effector proteins^3^. It was, therefore, initially expected that the phage-triggered defense mechanisms of Retron-Eco11 would differ from those of Retron-Eco1. However, with the exception of retron Ec48, the defense activity of which is overcome by phages λ and T7 through the use of inhibitors of the RecBCDE complex^2^, previous reports provided no conclusive evidence that a specific retron would respond to similar triggers from different phage types. Our findings support the notion that retrons providing defense against a broad spectrum of phages probably make use of the same mechanisms to sense and respond to phage infection.

UvsW is an ATP-dependent DNA helicase crucial for unwinding DNA structures, such as R-loops, during phage infection and DNA recombination. We hypothesized that the UvsW/D10 helicases might activate Retron-Eco11 by modifying the retron multicopy-single-stranded DNA (msDNA), an RNA-DNA hybrid molecule. This would imply that the phage triggers of the Retron-Eco11 system are core phage proteins capable of interacting with the msDNA, which is involved in DNA replication and recombination. Interestingly, the ATP hydrolysis-deficient mutant protein, UvsW-K141R, which lacks branched DNA-specific helicase activity, is not picked up by the retron system. This suggests that the Retron-Eco11 system may recognize helicase activity rather than the protein itself. The *E. coli* RecG helicase has similar functional properties to UvsW, but the absence of RecG does not lead to retron activation, implying that unique characteristics of UvsW/D10 helicases are central to this process. This specificity may also be related to the way in which these helicases interact with or access the retron complex.

Our findings demonstrate that Retron-Eco11, like other known retrons, forms a complex composed of three essential components: the RT, msDNA, and the PRTase effector. The Retron-Eco11 effector is not harmful on its own, but it may become activated on binding to the retron complex in response to UvsW/D10 phage helicase activity. Previous studies have indicated that the Ec86 retron effector interacts directly with the RT and msDNA, and that the msDNA wraps around the positively charged surface of the RT^5^. As for the Ec86 effector (nucleoside deoxyribosyltransferase-like), the predicted structure and architecture of the Retron-Eco11 effector suggest that it may have a two-lobed shape, with the N-lobe containing the core PRTase-like domain (α1-α14 and β1-β 9) and the C-lobe containing the winged-helix HTH DNA-binding domain (α15-α 21). It has also been suggested that the structure of Ec86 bound to its effector may represent the inactive state of this potentially toxic effector. It is, therefore, plausible that these two retron systems display essential functional and structural similarities.

As for Ec86, mutations of the catalytic active site of the RT and mutations of the branching G of the retron ncRNA abolished the defense activity of Retron-Eco11. Mutations affecting the branching G also disrupted the binding of the PRTase effector protein to the RT, demonstrating that msDNA is indispensable for the formation of the retron complex. Interestingly, it has been suggested that the long hairpin structure of the msDNA might play a role in phage recognition and interact with the effector protein, possibly inhibiting its activity^5^. The potential role of the DSLb hairpin structure of Retron-Eco11 and its flexible distal region in recognizing phage infections are therefore of interest.

The findings presented here provide compelling evidence for retron-mediated anti-phage defense that appear to operate through similar mechanisms, regardless of the phage type. These results pave the way for future studies to elucidate the mechanisms by which retrons sense and respond to phage infection.

## Acknowledgments

This work was supported by grant PID2020-113207GB-I00 from the MCIN/ AEI /10.13039/501100011033. We gratefully acknowledge José María del Arco Martín for his technical assistance.

## Authors’ contributions

FMGR performed the experimental work and contribute to writing-original draft. FMA, contribute to the identification and selection of retron systems for further analyses. MEW contributes to retron complex purifications assays. NTG, performed structural, statistical and phylogenetic analyses, writing-original draft its review and editing.

## Methods

### Bacteria and phage strains, plasmids and growth conditions

Phages T3, T5, T7, λ, M13, Qβ and ϕX174, and *E. coli* strains LE 392, Hfr 3000 U 432, MG1655, B, and BW25113 were acquired from the DSMZ-German Collection of Microorganisms and Cell Cultures GmbH. Phages T2, T4, P1, MS2 and ϕV-1 were obtained from the American Type Culture Collection (ATCC). *E. coli* strain MG1655 was used as host for phages T2, T4, T5, T7, P1, ϕV-1. For phages M13, MS2, Qβ and ϕX174, we used strain Hfr 3000 U 432 as the host. *E. coli* strain LE392 was used for assays of phage λ, and *E. coli* B was used for assays of phage T3. The antibiotics used were ampicillin (100 μg/ml) and chloramphenicol (200 μg/ml).

Retron-Eco11 was produced by GenScript Corp. and provided cloned into pUC57mini to generate the p57mA3 plasmid, which confers ampicillin resistance. Mutants of the PRTase, branching G and RT were also obtained from GenScript Corp. and were delivered cloned into pUC57mini as p57mA3_PRTase, p57mA3_C and p57mA3_RT, respectively. RT-Eco11 was C-terminally tagged with 3xFlag tag with the NEBuilder HiFi DNA Assembly kit from New England BioLabs Inc. (NEB). The p57mA3 plasmid was digested with MscI restriction enzyme and the following oligomers were used: 3C-3xFlag-5’ (5’-GTAAATATAATGAGATTAATTGGCCATTTCTAGAGGTTCTCTTTCAAGGGCCAGCAGACTACAAAGACCATGACGGTGATTATAAAGATCATGACATCGATTACAAGGATGACGATGACAAGTGATTTTTGGGTGTTGTATAAGTG-3’) and 3C-3xFlag-3’ (5’-CACTTATACAACACCCAAAAATCACTTGTCATCGTCATCCTTGTAATCGATGTCATGATCTTTATAATCACCGTCATGGTCTTTGTAGTCTGCTGGCCCTTGAAA GAGAACCTCTAGAAATGGCCAATTAATCTCATTATATTTAC-3’). Cloning was performed according to manufacturer’s instructions to generate p57mA3-RTCF. The same strategy was used to construct p57mA3mRTCF, which carries a modified form of the YSDD motif of the RT (YSAA) and a 3xFlag tag at the C-terminal end of the RT. We digested p57mA3-RT with MscI and the same oligomers, 3C-3xFlag-5’ and 3C-3xFlag-3’, were used. For complex purification, the Retron-Eco11 operon was amplified from p57mA3-RTCF with primers 3A3_PRTase_F (5’ TGTTTAACTTTAAGAAGGAGATATACATGAGCTCTCATGAAATAACATTAG ATAAG-3’) and 3C-RT_R (5’ TGCGGATGGCTCCAGCTAGCTGCTGGCCCTTGAAAGAGAA-3’) and subcloned using NEBuilder HiFi DNA Assembly into a modified pET45b (+) vector such that the RT contained a C-terminal TwinStrep tag (WSHPQFEKGGGSGGGSGGSAWSHPQFEK), generating pMW261.

Phage helicases UvsW and D10 were amplified by PCR and inserted into pBAD33 under the control of an arabinose-inducible promoter, to create the p33UvsWwt and p33D10wt plasmids. The *uvsW* gene was amplified with primers uvsW5’ (5’-CTGGTACCATGGATATTAAAGTACATTTTCACG-3’) and uvsW3’ (5’-CTGTCGACTTAAAAGCTTTCTTCTACTTCATC-3’) with total DNA extracted from T4 phage as the template. The D10 helicase was amplified from total DNA isolated from T5, with the primers D10-5’ (5’-CTGGTACCATGAAGGTTGTTATATCTAATAAAG-3’) and D10-3’ (5’-CTGTCGACTTATGAGCTGTTGCCAAATGC-3’). For p33UvsWmut6, the mutant *uvsW* gene was amplified with primers uvsW5’ and uvsW3’ from total DNA isolated from the T4 mutant 6 phage, and was inserted into pBAD33. The p33D10mut13 plasmid was constructed in a similar manner, with primers D10-5’ and D10-3’ used for amplification of the mutant D10 gene from T5 mutant 13 phage DNA. The K141R substitution of UvsW was generated by overlapping PCR fragments obtained with the oligomers uvsW5’ and K141R Rv (5’-TGAATTAAAGATCTACCTGCAGATG-3’) for the first PCR and uvsW3’S (5’-CAAGCTTGCATGCCTGCAGG-3’) and K141R Fw (5’-CATCTGCAGGTAGATCTTTAATTCA-3’) for the second PCR. For both PCRs p33uvsW was used as the template. A third PCR was performed with the uvsW5’ and 3’uvsWS primers, with 1 μl of the first and second PCRs used as the template. The PRTase gene was amplified with the oligomers PRTaseWH Fw (5’-CTGGTACCATGAGCTCTCATGAAATAACATT-3’) and PRTaseWH Rv (5’-CTGTCGACTTATGGCCTCCTCAACGTTG-3’), with p57mA3 as the template. The PCR product was sequenced and inserted between the KpnI and SalI sites of pBAD30 to yield p30PRTase.

### msDNA isolation

We isolated msDNA from FLAG immunoprecipitates as follows: 50 μl of immunoprecipitated was subjected to extraction with phenol:chloroform:isoamyl-alcohol (25:24:1, pH 8) and nucleic acids were precipitated by incubation overnight at −80°C with 2 volumes of 100% ethanol and 1/10 volume of 3 M sodium acetate pH 5.2. The precipitated nucleic acids were obtained by centrifugation at14000 rpm at 4°C for 60 min. Pellets were air-dried and resuspended in 10 μl of distilled water containing RNase A/T1 and the suspension was then incubated at 37°C for 30 min. Samples were stored at −80°C until use. For msDNA visualization, samples were mixed with TBE loading buffer and incubated at 75°C for 5 min and then cooled on ice for 5 min. Samples were loaded onto a 10% denaturing polyacrylamide gel and run in 1xTBE for 90 min at 20 W. Gels were stained by incubation for 15 min in a GelRed bath.

### Identification of the 3’ end of retron Eco11

The msdDNA cDNA was isolated from acrylamide gels and 100 ng was used to extend the 3’ end with dCTP or dGTP and terminal deoxynucleotidyl transferase (TdT) from NEB, according to the manufacturer’s instructions. For each tailing reaction, 5 μl of the reaction mixture was subjected to PCR amplification with the oligomers A3-3’ (5’-CGAAGGGGTAAGTAAAATCG-3’) and PoliGH (5’-GGGGGGGGGGGGH-3’) in the presence of dCTP or A3-3’ and PoliCD (5’-CCCCCCCCCCCCD-3’) in the presence of dGTP. The PCR products were gel-purified, inserted into pGEM-T and 10 clones from each tailing reaction were sequenced to characterize the 3’ end.

### Phage plaque assays

Double-layer plaque assays^30^ were performed. *E. coli* host strains were grown to saturation at 37°C in LB. We then added 200 μl of the *E. coli* culture to 10 ml top agar (LB, 0.5% agar, 5 mM MgSO_4_ and the appropriate antibiotic) and poured the mixture onto 12 x12 cm LB-agar plates containing 5 mM MgSO_4_ and the appropriate antibiotic. We spotted 10 μl of ten-fold dilutions of phages in liquid LB onto the plates and plaques were counted after overnight incubation at 37°C. If individual plaques were too small to be counted, the most concentrated dilution at which no plaques were visible was recorded as having a single plaque.

### Isolation and amplification of mutant phages

For the isolation of mutant phages capable of escaping Retron-Eco11 defenses, we plated phages on a lawn of MG1655 bacteria expressing the Eco11 defense system (p57mA3) in a double-layer plaque assay. We picked five individual plaques into 100 μl LB, which was left at room temperature for 1 h, with mixing several times by vortexing to release the plaques from the agar. The phages were centrifuged at 3200 x *g* for 10 min to remove agar and bacteria and the supernatant was transferred to a new tube. The ability of the collected phages to escape the Eco11 defense system was assessed in plaque assays with a strain expressing the defense system and a strain lacking the defense system (negative control). Ten-fold serial dilutions were performed for the ancestor phage (WT phage used for the original double-layer plaque assay) and the phages collected from plaques formed on the strain with the defense system. Isolated phages against which defenses were weaker than for the ancestor phage were further propagated by picking a single plaque formed on cells expressing Retron-Eco11 into a 10 ml liquid culture of Eco 11 cells grown in LB plus 5 mM MgSO_4_ at an OD_600_ of 0.3. The culture was incubated for 6 h at 37°C with shaking at 175 rpm. The lysate was then centrifuged at 3200 x *g* for 10 min and the supernatant was passed through a filter with 0.2 μm pores.

### Phage genome sequencing and analysis of phage mutants

Phage DNA was extracted from 500 μl of a high-titer phage lysate (>10^7^ PFU/ml). Samples were treated with DNAse-I (20 μg/ml) and incubated for 1 h. DNA was extracted with the Qiagen DNeasy blood and tissue kit with a protocol beginning with proteinase K treatment. Libraries were prepared for Illumina sequencing. Reads were mapped onto phage reference genomes (GenBank accession numbers: NC_000866 (T4 phage) and NC_005859 (T5 phage)) with Geneious Prime software. Only mutations that occurred in the isolated mutants but not in the ancestor phage were considered.

### Growth curves

*E. coli* MG1655 harboring the retron sytem (p57mA3) and the UvsW, D10 phage helicases or their mutant forms UvsWK141R or D10mut13 were grown overnight in LB supplemented with ampicillin, chloramphenicol and 1% glucose. We then used 2 ml of each culture to inoculate each of two flasks containing 50 ml LB plus ampicillin and chloramphenicol. The flasks were incubated at 37°C until an OD_600_ of 0.3 was reached. Arabinose was then added, at a final concentration of 0.2%, to one of the two flasks. The flasks were incubated for another four hours at 37°C. Samples were taken from each of the flasks at 0 h (just before the addition of arabinose to one of the flasks), and 2 h and 4 h after arabinose addition. The samples were diluted and plated on LB-agar plus ampicillin and chloramphenicol plates for the counting of viable cells.

### Spot growth tests

*E. coli* MG1655 harboring the retron sytem (p57mA3) or a negative control (p57m, empty plasmid) along with an arabinose-inducible plasmid containing *uvsW* (p33uvsW), the D10 gene (p33D10) or an empty plasmid (pBAD33) was grown overnight in LB supplemented with ampicillin, chloramphenicol and 1% glucose. The cultures were then subjected to a 10-fold serial dilution, and 10 μl of each dilution was spotted onto an LB agar plates supplemented with antibiotics, with or without 0.2% arabinose. Plates were incubated at 28°C overnight. The spot tests shown in Figure 6A were performed as described above but with *E. coli* BW25113 harboring p30PRTase, in the presence of ampicillin as the only antibiotic.

### Affinity purification

We inoculated 200 ml LB supplemented with the appropriate antibiotics with 2 ml of an overnight culture of *E. coli* MG1655 harboring plasmids expressing Flag-tagged RT or the negative control (the same plasmid with no Flag-tagged gene). Cultures were incubated at 37°C until an OD_600_ of 1.2 was reached. They were then centrifuged at 12000 x *g* for 10 min at 4°C and the supernatant was discarded. The pellet was washed with 50 ml PBS and resuspended in 8 ml of lysis buffer (50 mM Tris-HCl pH 7.4, 150 mM NaCl, 1 mM EDTA and 1% TRITON X-100) supplemented with protease inhibitors. Samples were sonicated with four pulses of 15 s each and were then centrifuged at 12000 x *g* for 10 min at 4°C. The pellet was discarded and 100 μl Anti-FLAG M2-Agarose Affinity Gel (prepared according to the manufacturer’s instructions) was added to the supernatant, which was then incubated overnight at 4°C with gentle shaking. Samples were centrifuged for 30 seconds at 8200 x *g* and the supernatant was discarded. The precipitated resin was either used for co-incubation experiments (see below) or processed to isolate components of the retron complex. For retron complex analysis, the resin was washed three times with 0.5 ml wash buffer (50 mM Tris-HCl, pH 7.4, 150 mM NaCl). For elution, the resin was incubated with 100 μl wash buffer supplemented with 150 ng/μl 3X Flag peptide for 30 min at 4°C with gentle shaking, and then centrifuged at 8200 x *g* for 30 s. The supernatant containing the proteins was stored at −20°C.

### Retron complex purification

The retron complex was overexpressed in BL21(DE3) from plasmid pMW261 in autoinduction media containing ampicillin. Cells (2L) were grown at 37°C for 8 h then at 18°C for 16 h. For coexpression with UvsW, BL21(DE3) was co-transformed with pMW261 and either p33UvsWwt or p33UvsWK141R and overnight cultures were grown in the presence of ampicillin, chloramphenicol, and 0.2% glucose. 10 mL of culture was used to inoculate 1 L of LB containing ampicillin, chloramphenicol and 2 mM MgCl_2_ and cells were grown at 37°C until OD600 reached 0.6-0.8, at which point 0.5 mM IPTG was added to induce retron expression. After 2 h at 37°C, 0.1% arabinose was added to induce UvsW expression, and after a further 2 h cells were harvested and frozen at −80°C. Retron complexes were purified from all three conditions identically: cells were suspended in lysis buffer (50 mM Tris-HCl pH 74, 500 mM NaCl, 5% glycerol, 5 mM beta-mercaptoethanol) and lysed in a LM20 microfluidizer (Microfluidics). Cleared lysate was bound to Strep-Tactin Superflow Plus resin (Qiagen). The resin was washed first with lysis buffer and then with RNP buffer (20 mM HEPES-KOH pH 7.9, 250 mM potassium acetate, 2.5 mM magnesium acetate) before elution in RNP buffer containing 5 mM desthiobiotin. Protein-containing fractions were pooled and concentrated in a Vivaspin 20 centrifugal concentrator (50000 MWCO; Sartorius). Purified complexes were analyzed by spectrophotometry (Nanodrop) and all had A260/A280 ratios of 1.8, indicative of copurifying nucleic acids. The concentrated fractions were extracted with phenol:chloroform:isoamyl alcohol (25:24:1), precipitated with ethanol and suspended in RNase-free water. 200 ng of nucleic acids from each complex were mixed in 1xDNase I buffer (NEB) with either RNase-free DNase I (NEB), 100 ng of RNase A, or a no enzyme control, and incubated at 37°C for 30 min. Reactions were boiled at 95°C for 150 s, placed on ice, and run on a precast 10% acrylamide TBE-Urea gel (Invitrogen) at 400 V using 1xTBE running buffer preheated to ∼50°C. The gel was stained with SYBR Gold (Invitrogen) in 1xTBE for 15 min and visualized using a ChemiDoc (Bio-Rad).

### Co-incubation experiments

We inoculated 200 ml LB plus chloramphenicol with 2 ml of an overnight culture of *E. coli* BW25113 harboring p33uvsW or p33uvsD10. The cells were cultured at 37°C until an OD_600_ of 0.3 was reached. Arabinose was added to a final concentration of 0.2% and culture was continued until an OD_600_ of 1.2 was reached. Cultures were centrifuged at 12000 x *g* for 10 min at 4°C and the supernatant was discarded. The pellet was washed with 50 ml PBS and resuspended in 8 ml lysis buffer (50 mM Tris-HCl pH 7.4, 150 mM NaCl, 1 mM EDTA and 1% Triton X-100) supplemented with protease inhibitors. Samples were sonicated with four pulses of 15 s each and centrifuged at 12000 x *g* for 10 min at 4°C. The pellet was discarded and the supernatant was mixed with the resin obtained as described in the affinity purification section (see above) and incubated overnight at 4°C with gentle shaking. Samples were then centrifuged for 30 seconds at 8200 x *g* and the supernatant was discarded. The resin was washed three times with 0.5 ml wash buffer (50 mM Tris-HCl, pH 7.4, 150 mM NaCl). For elution, the resin was incubated with 100 μl wash buffer supplemented with 150 ng/μl 3X Flag peptide for 30 min at 4°C with gentle shaking, and then centrifuged at 8200 x *g* for 30 s. The supernatant containing the proteins was stored at −20°C

### Proteomic analysis of affinity purification

Proteomic analysis was performed at the Proteomics Service of *Instituto de Parasitología y Biomedicina “López-Neyra”* (CSIC, Granada). A volume of supernatant from affinity purification containing 1.5 μg of protein was subjected to electrophoresis for 10 min in a 4% SDS-PAGE gel, which was then stained with Coomassie blue. The band was excised and subjected to in-gel tryptic manual digestion for 18 h a 30°C. Peptides were extracted from the gel in 0.2% trifluoroacetic acid (TFA), 30% acetonitrile. Samples were dried and stored at −20°C until use. Before nLC-MS/MS analysis, samples were resuspended in 0.1% FA, 0.3% acetonitrile. The peptides were fractionated on an Easy n-LC II chromatography system (Proxeon) connected to an Amazon Speed ETD mass spectrometer (Bruker Daltonics). Chromatography was performed with a C18 reverse-phase analytical column (15 μm x 15 cm, 2.6 μm, 100 A) over a 180 min organic gradient from 5 to 30% B (buffer A: 0.1% FA in H_2_O; buffer B: 0.1% FA in acetonitrile) with a flow rate of 300 nl min^−1^. The ion trap was set to analyze the survey scans in the mass range m/z 390 to 1400. Search criteria included carbamidomethylation of cysteine as a fixed modification and oxidation of methionine as a variable modification. The fold-enrichment of pull-down proteins was calculated, and statistical significance was evaluated with R.

### Phylogenetic tree construction

We used MAFFT software^31^ and progressive methods for MSAs. The unrooted tree was constructed from an alignment of 2564 sequences carrying the phosphoribosyltransferase domain (IPR000836) and 127 sequences from the retron type III-A associated protein cluster 67_3^3^. It was generated with the FastTree program^32^ with the WAG evolutionary model and the discrete gamma model, with 20 rate categories.

## RESOURCE AVAILABILITY

### Availability of materials

Bacterial and viral strains generated in this study are available upon request.

---

**Figure S1.**
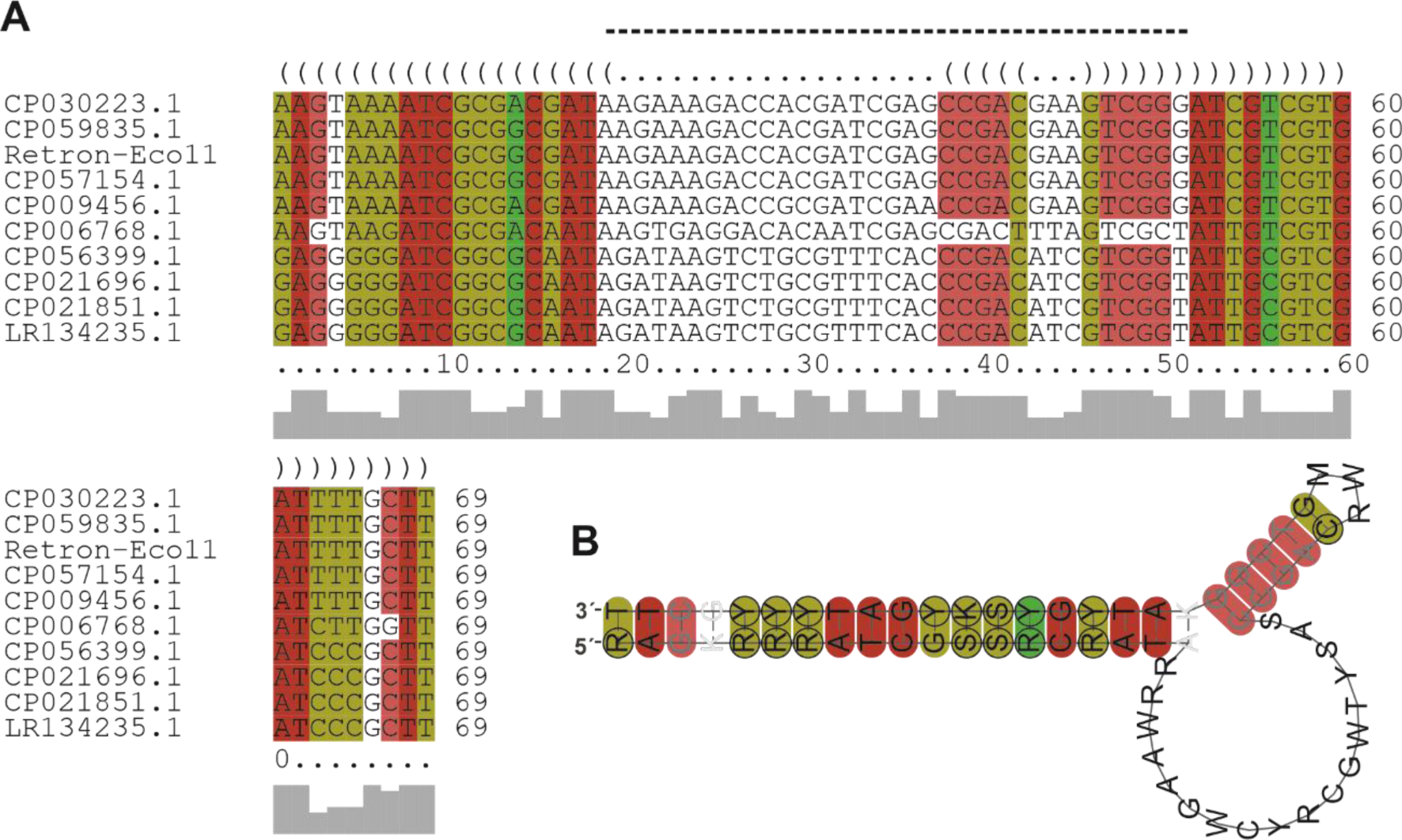
Structural alignment corresponding to the DSLb stem loop from type III-A retrons. (**A**) Alignment of DSLb DNA nucleotide sequences showing 66% pairwise identity, as performed with Locarna^36^. Conserved nucleotides are indicated, and the structurally flexible DSLb hairpin loop is marked by a dashed line above the sequence. CP006768.1:2313606-2313848_(reversed) *Acinetobacter baumannii* ZW85-1, CP059835.1:3020050-3020289 *Escherichia coli* strain tcmA_3, CP009456.1:1763831-1764070_(reversed) *Yersinia enterocolitica* strain FORC-002, CP056399.1:4400487-4400716_(reversed) *Citrobacter* sp. RHBSTW-00570, CP021696.1:4757031-4757260_(reversed) *Klebsiella pneumoniae* strain AR_0158, LR134235.1:1507974-1508193 *Klebsiella variicola* strain NCTC9668 genome assembly, chromosome: 1, CP057154.1:5085011-5085250_(reversed) *Escherichia coli* strain RHB35-C09, CP021851.1:1088677-1088896 *Enterobacter cloacae* strain A1137, and CP030223.1:4679774-4680013_(reversed) *Salmonella enterica* strain SA20083039. (**B**) The consensus structure of the alignment as predicted by RNAalifold.

**Figure S2.**
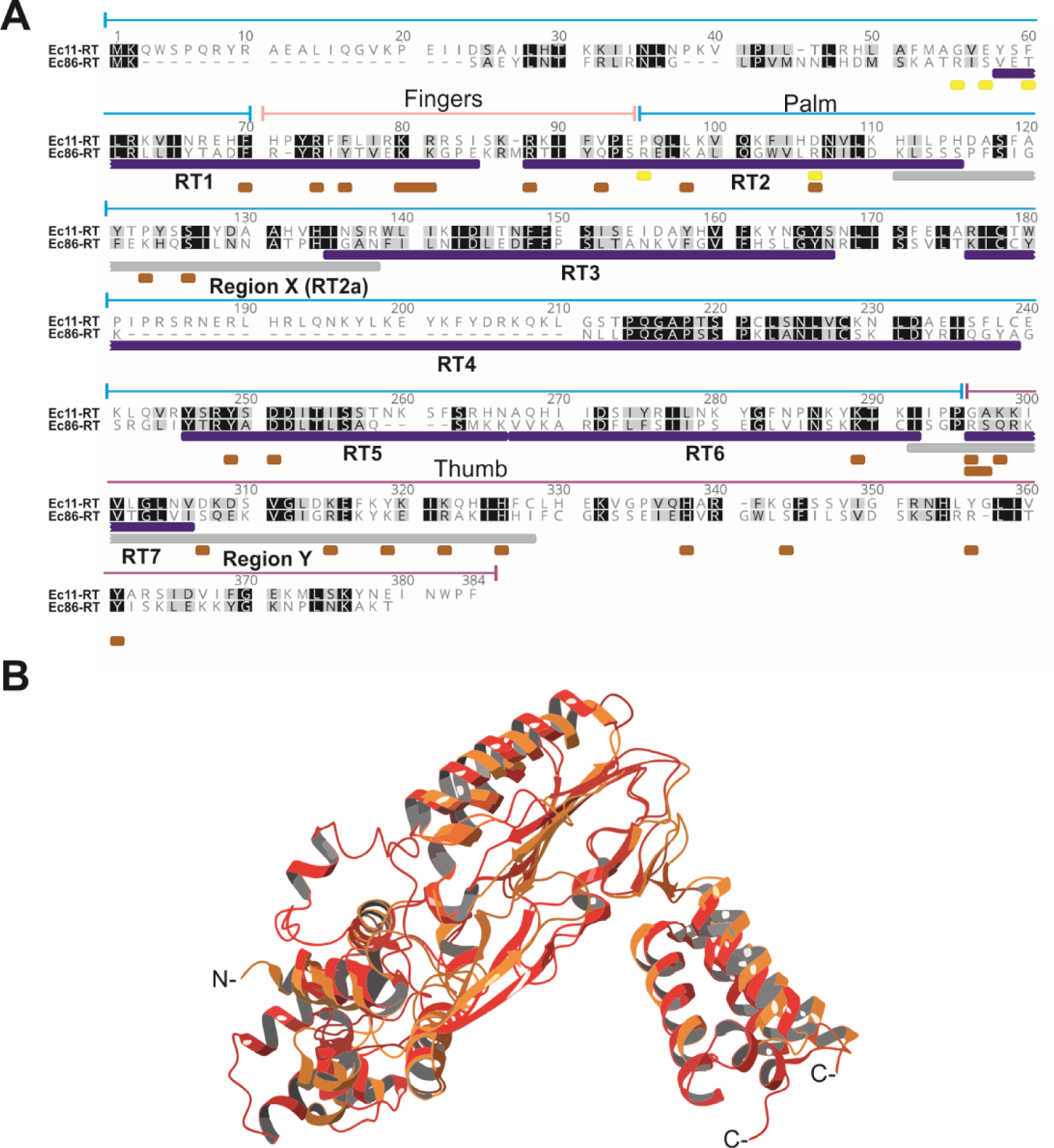
Retron-Eco11 reverse transcriptase. (**A**) Alignment of the reverse transcriptases encoded by the Eco11 and Ec86 retrons. Identical and similar amino acids are shaded in black and gray, respectively. The RT-Eco11 aligned with the RT-Eco1, showing a pairwise identity of 24.5%, encompassing the RT1-7 regions and the retron-specific regions, denoted X and Y. In these retron-specific regions, the conserved NAxxH motif of Ec86 is replaced by AAxxH, and the VTG is replaced by VLG. Ec86 residues involved in RT-msDNA interaction are indicated by a brown bar below the sequence whereas residues involved in interactions with the effector protein are indicated by yellow bars^5^. Palm domain, cyan; finger domain, pink; thumb domain, purple. (**B**) Superimposition of the predicted 3D structure of RT-Eco11 (in red) generated by AlphaFold2 on that of the RT-Eco1 (in orange) (Ec86 complex, Protein Data Bank (PDB): 9V9U). The superimposition yielded a root mean square deviation (RMSD) of 2.55 over 320 Cα atoms. N and C denote the N and C-termini, respectively, of the proteins

**Figure S3.**
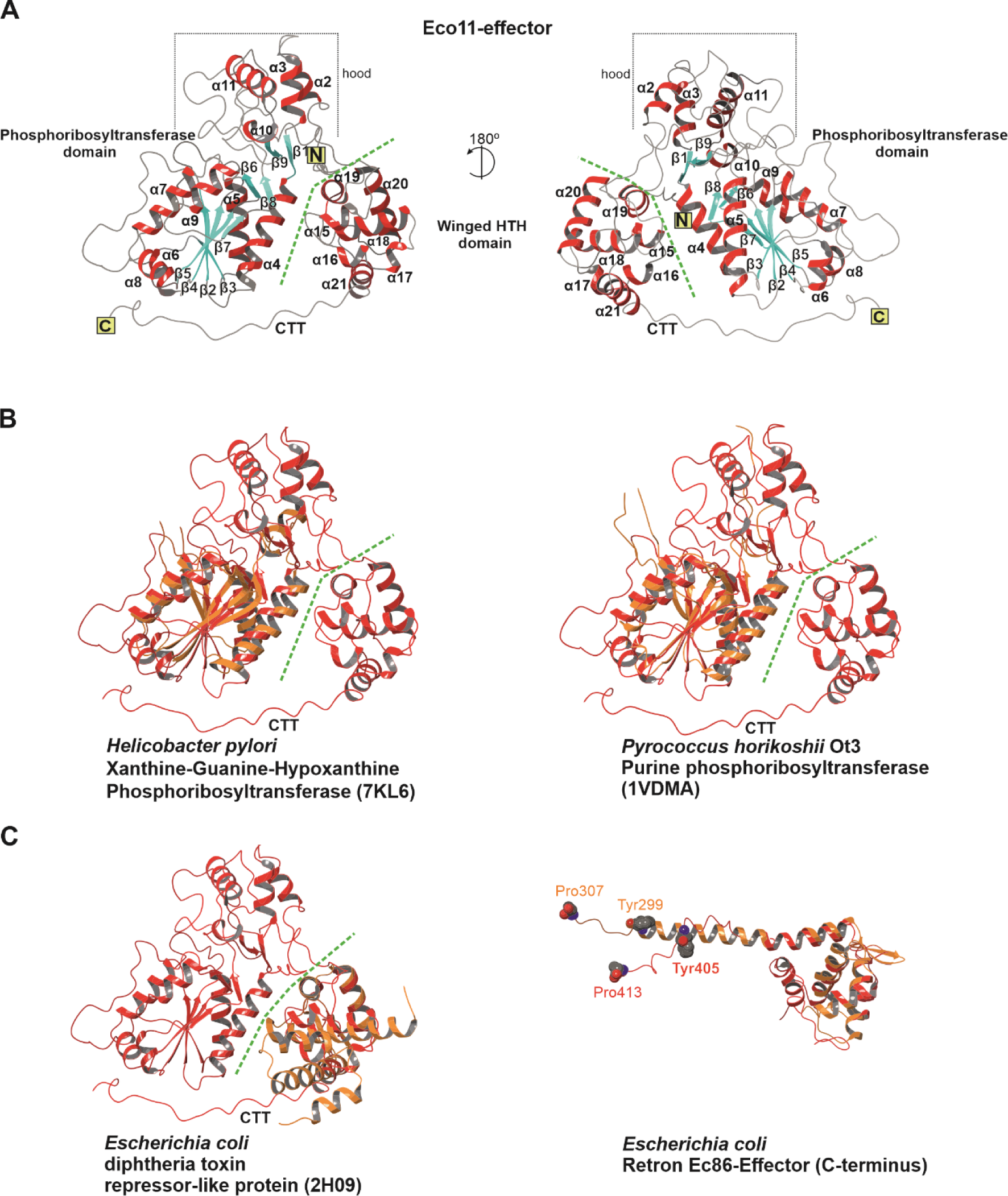
Insights into the structural characteristics and relationships of the Eco11-effector PRTase. (**A**) Predicted 3D structure of the Eco11-effector PRTase according to AlphaFold 2. The phosphoribosyltransferase and winged-HTH domains are labeled and separated by a dashed green line. Like other PRTases, the Eco11 effector possesses a subdomain called the “hood” at the C-terminal edge of the core B-sheet. This hood consists predominantly of residues located at the N-terminus of the protein^14^. (**B**) Superimposition of the predicted 3D structure of the Eco11 PRTase (highlighted in red) on the structures of the xanthine-guanine-hypoxanthine PRTase (PDB: KL6) from *H. pylori* (with an RMSD of 3.07 over 130 Cα atoms) and the purine PRTase (PDB:1VDMA) from *P. horikoschii* (with an RMSD of 3.09 over 132 Cα atoms). (**C**) Superimposition of the predicted 3D structures of Eco11 PRTase (in red) in its winged-HTH domain with the *E. coli* diphtheria toxin (PDB: 2H09) repressor-like protein (RMSD of 2.00 over 72 Cα atoms), and the effector-bound Ec86 complex (PDB: 7XJG) (RMSD of 2.55 over 73 equivalent positions). The latter has a structurally similar terminal peptide, flanked by tyrosine (residue 299) and proline (residue 307) residues equivalent to the Tyr405 and Pro413 residues of the Eco11 effector.

**Figure S4.**
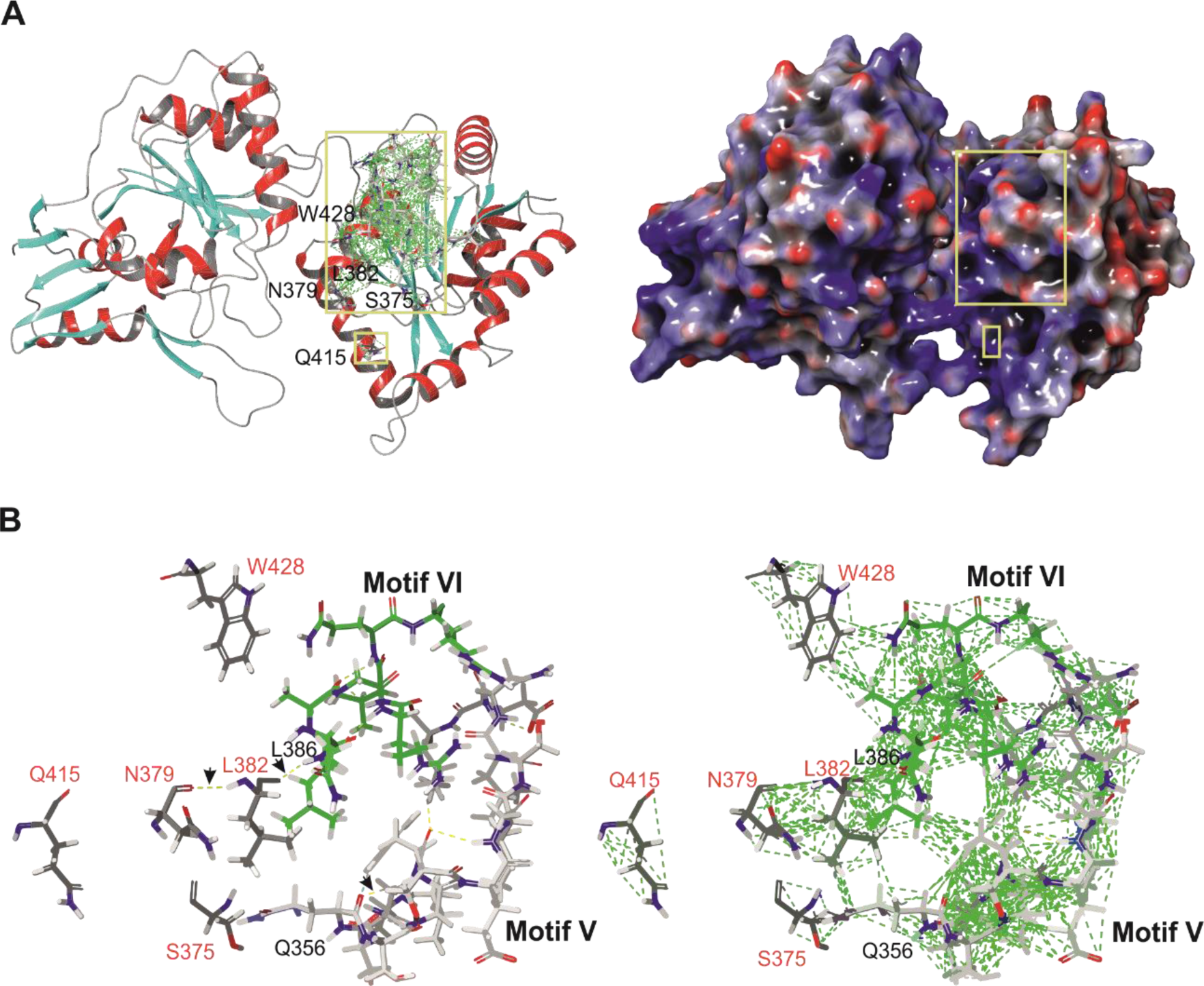
Structural features of phage T5 D10 amino-acid substitutions. (**A**) Predicted structure of phage T5 D10 helicase obtained with AlphaFold2, with the residues substituted in the T5 phage escape mutants highlighted. The electrostatic surface of the protein is displayed on the right side of the panel. Blue and red surfaces represent positive and negative potentials, respectively. The locations of the residues mutated in the T5 phage escape mutants are highlighted. (**B**) Detailed enlargement of the structural regions of the D10 protein encompassing motif V (highlighted in gray) and VI (highlighted in green), which interact with the residues substituted in the phage escape mutants. Hydrogen bonds presumed to be important (indicated by yellow dashed lines) between these residues and residues in motifs V and VI are indicated by arrows.

**Figure S5.**
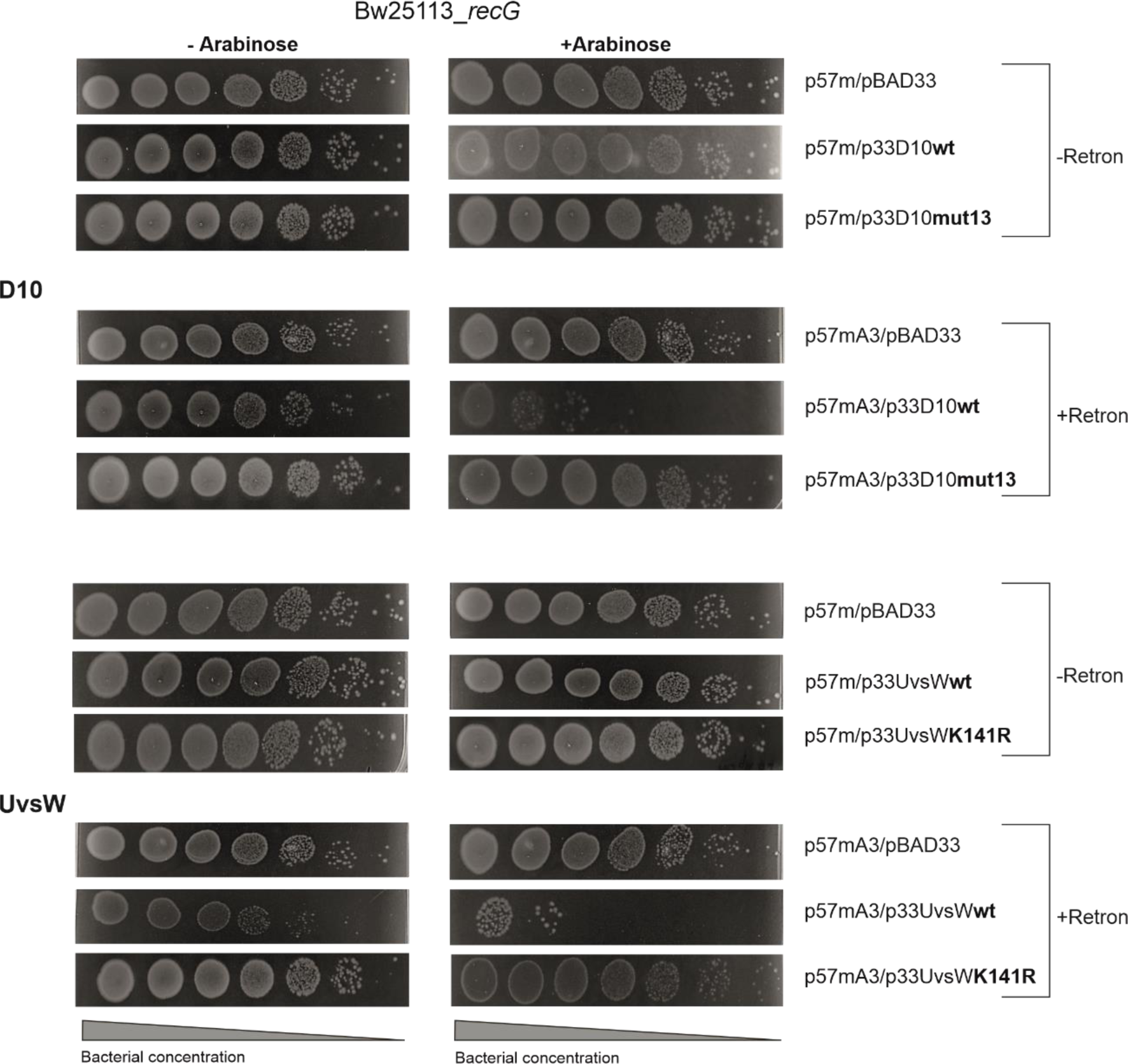
Expression of the Retron-Eco11 system and the T4 UvsW and T5 D10 helicases in an *E. coli* Δ*recG* mutant. The Δ*recG* mutant strain of *E. coli*, BW25113, was used in the experiments. Plasmid constructs, bacterial viability measurements, and experimental procedures are as described in Figure 6.

